# Dose-dependent power and connectivity modulation of low frequency oscillations through transcranial magnetic stimulation in non-human primates

**DOI:** 10.64898/2026.01.13.699238

**Authors:** Malte R. Güth, Nipun D. Perera, Gary Linn, Kurt Masiello, Brent Butler, Brian E. Russ, Charles E. Schroeder, Arnaud Falchier, Alexander Opitz

## Abstract

Transcranial magnetic stimulation (TMS) is a powerful non-invasive tool for safely modulating neural activity in humans. In particular, the left dorsolateral prefrontal cortex (DLPFC) is a common target site for clinical interventions in disorders such as treatment-resistant depression. Yet, clinical trials investigating the efficacy of TMS often lack neural markers of target engagement of the DLPFC. Local field potentials (LFPs), such as prefrontal theta oscillations, have been implicated in the clinical symptoms of these disorders. However, non-invasive electroencephalography (EEG) recordings in humans are limited by their spatial resolution and challenges of interpreting EEG signals. In this study, we investigate the effects of single-pulse TMS applied to the left prefrontal cortex in non-human primates on LFPs recorded through intracranial EEG. Compared to sham TMS, the intensity of active TMS pulses scaled with LFP power changes in a 1-13 Hz range at contacts close to the stimulation site in the prefrontal cortex (e.g., caudate nucleus, anterior cingulate cortex, insular cortex) as well as contacts that were more distal (e.g., posterior cingulate cortex, temporal lobe). To test how TMS modulates connectivity between these regions, we conducted a phase-based connectivity analysis. TMS pulses initially enhanced and then disrupted connectivity at 1-13 Hz between the stimulation site and other contacts. Connectivity rebounded approximately 1500 ms post-stimulation. Only the initial enhancement in connectivity scaled with TMS intensity. Our results demonstrate a dose-dependent power modulation of low frequency LFPs across prefrontal, parietal and temporal cortical regions by single pulses. Furthermore, they show that TMS applied over the left prefrontal cortex can enhance and interrupt short- and long-range connectivity. Our study advances the understanding of the effects of TMS on brain oscillations and connectivity with direct relevance for clinical applications in neuromodulation therapies.

## Introduction

Transcranial magnetic stimulation (TMS) is a crucial intervention tool in research and clinics, capable of non-invasively exciting or disrupting neural activity with millisecond precision (1). TMS is used to probe brain function and connectivity using its ability to modulate cortical excitability in a targeted and temporary manner. For example, repetitive TMS (rTMS) applied over the dorsolateral prefrontal cortex (DLPFC) has been shown to be effective as an intervention in several psychiatric disorders, in particular depression (2–4). Theta burst stimulation (TBS) as well as 10-Hz and 20-Hz protocols targeting the left DLPFC have generally demonstrated anti-depressant effects (5–7), most notably in treatment-resistant depression (8–11). Magnetic resonance imaging (MRI) studies have highlighted the DLPFC as a key structure for working memory (12,13), executive function (14,15), cognitive control (16,17), decision-making (18–20), and emotion regulation (21–23). In individuals with depression, the DLPFC exhibits both morphological abnormalities, such as a reduction in gray matter volume (24) as well as functional abnormalities, such as an increased sensitivity to negative emotional stimuli (25). Given the DLPFC’s critical role in these cognitive and emotional processes, the DLPFC has become a key target for neuromodulation therapies. The DLPFC has also been targeted due to its direct connections to deeper brain regions, such as the anterior cingulate cortex (26–28) and adjacent cortical regions that are part of a larger cortical network (i.e., parietal lobe) (29–31).

However, while the prefrontal cortex (PFC) is commonly targeted with TMS, it does not provide direct measures of engagement through TMS, hindering the evaluation of treatment protocols and targeting procedures. The primary motor cortex owes its status as a benchmark TMS targeting site to the availability of motor evoked potentials, which serve as a direct readout of cortical excitation. Prefrontal cortical regions do not offer an equivalent observable output from TMS application, but experimentally evoked event-related potentials (ERPs), evoked oscillatory activity, and TMS evoked potentials (TEPs) recorded via electroencephalography (EEG) have been suggested as markers for TMS treatment response (32–34). These EEG metrics provide a valuable but indirect measure of the brain’s oscillatory activity. Midfrontal theta (4-8 Hz) and alpha (9-12 Hz) oscillations in particular have been connected to numerous functions attributed to DLPFC activation as a whole, such as exerting cognitive control (35,36), decision-making (37,38), working memory (Griesmayr et al., 2014; Popov et al., 2018; Riddle et al., 2020; Scheeringa et al., 2009), reinforcement learning and feedback processing (43–45). While several studies have tested the effects of TMS on these oscillations (e.g., Chung et al., 2018; Riddle et al., 2020), the limitations of non-invasive EEG in terms of spatial resolution hinder a comprehensive understanding of how TMS influences these oscillatory patterns at a local and network level. For example, when investigating sensor-to-sensor connectivity using phase-based measures of EEG, the high degree of smoothing between sensors due to volume conduction limits the validity of connectivity estimates (47).

In non-human primates (NHPs), intracranial electroencephalography (iEEG) offers a more direct and precise method for measuring neural activity, enabling researchers to explore the effects of TMS with greater spatial specificity in both the targeted and distal regions (48,49). NHP models have been leveraged in the past to study the effects of single-pulse TMS (spTMS) on single-cell activity. spTMS elicited transient increased spiking in cortical areas beneath the coil within a few milliseconds of stimulation (50,51), followed by a reduction in spiking activity and a return to baseline. In terms of macro-scale activity, we previously demonstrated a dose-dependent relationship between the intensity of spTMS and the amplitude of early TEPs (i.e., N50) assessed using iEEG in NHPs (52). We observed a sigmoidal relationship between increases in TMS intensity and increases in the amplitude of the N50. While this previous study focused on TMS-evoked potentials, here we investigate the effects of single biphasic TMS pulses in the left PFC on spectral power and network connectivity across prefrontal, parietal, and temporal regions. Specifically, we investigated how varying intensities of TMS pulses modulate the power of LFPs. Moreover, we explore the pulse-locked changes in phase-based connectivity between cortical regions, with a focus on the connectivity between the stimulation site and the remaining contacts. Our work provides novel insights into the immediate neural effects of TMS on oscillatory brain activity and connectivity in NHPs in a target region directly relevant for clinical applications in humans.

## Methods

### Subjects and surgical procedures

All experimental protocols were approved by the Animal Care and Use Committee at the Nathan S. Kline Institute for Psychiatric Research. The study involved two adult rhesus macaques (Macaca mulatta): Monkey W (female, 5 kg) and Monkey H (male, 10 kg). Both monkeys were implanted with an MRI-compatible polyetheretherketone headpost, positioned over the occipitoparietal region. Three stereo EEG (sEEG) depth electrodes (Monkey H: 29 contacts, Monkey W: 28 contacts), with 5 mm spacing (Ad-Tech), were permanently placed through a skull opening over the left occipital cortex. The electrodes were oriented along a posterior-to-anterior axis targeted at the medial prefrontal cortex, frontal eye field, and auditory cortex. For Monkey W the third sEEG electrode was inserted in the superior temporal cortex and for Monkey H in the medial and inferior temporal cortex (Figure 1A).

**Figure 1.**
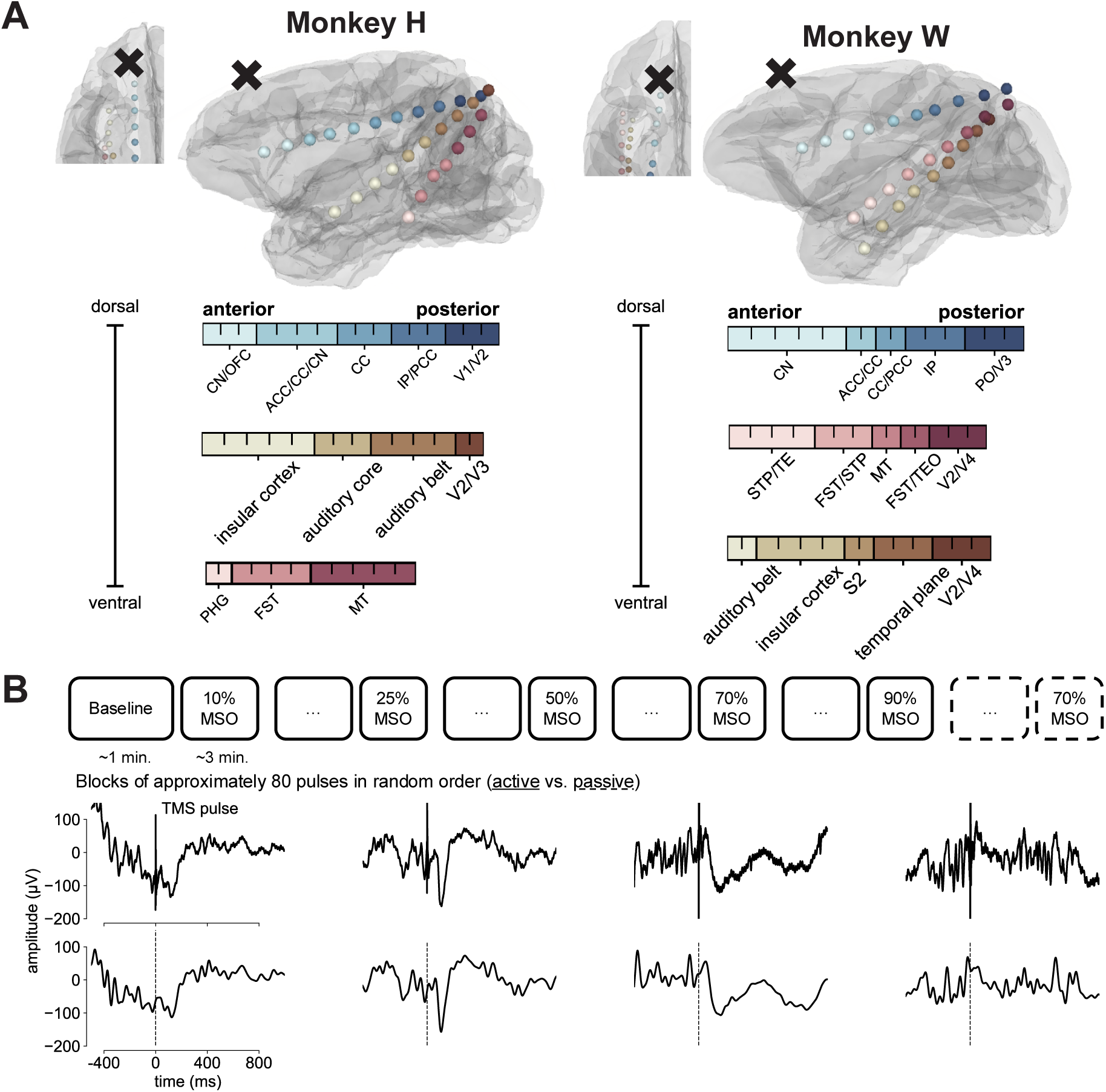
**A**: Overview of recording electrodes (colored spheres) and TMS coil locations (marked by **x**) on each monkey’s reconstructed gray matter (GM) surface. The three stereo-electrodes are depicted with distinct colors (blue, yellow, red). Individual spheres represent contact locations within an electrode. Contact colors are scaled according to the anterior to posterior position. Inset at top left shows an axial view of the left frontal GM. Color bars list the anatomical regions closest to each contact, with each tick inside a color bar representing one contact. CN: caudate nucleus, OFC: orbitofrontal cortex, CC: corpus callosum, ACC: anterior cingulate cortex, PCC: posterior cingulate cortex, IP: intraparietal area, V1: visual area 1, V2: visual area 2, V3: visual area 3, PHG: parahippocampal gyrus, FST: fundus of the superior temporal visual area, MT: middle temporal area, PO: preoptic area, TE: anterior inferotemporal area, STP: superior temporal plane, TEO: Temporal Extrastriate Visual Cortex, V4: visual area 4, S2: secondary somatosensory cortex. **B**: Top row illustrates the timeline of the experimental procedure with alternating TMS conditions (solid lines: active, dashed: passive). Middle and bottom rows show single-trial examples for 70% MSO stimulations at anterior contacts for Monkey H and Monkey W. Middle row shows the raw trace and bottom row shows the same trace after full preprocessing, including artifact interpolation. Dashed vertical lines mark the timing of TMS pulses.

### Magnetic resonance imaging and electrode localization

MRI scans for both monkeys were acquired using a Siemens TIM Trio 3T scanner. These included T1 and T2 spin echo sequences with the following parameters: voxel size = 0.5 × 0.5 × 0.5 mm; flip angle = 80°; TR = 2600 ms; TE = 3.55 ms; and TI = 900 ms. Electrodes were localized on post-implantation MR images, which were registered to pre-implantation scans and aligned with a standard stereotaxic atlas (53) using the FSL FLIRT registration tool (54–56). An expert in nonhuman primate anatomy identified the brain structure closest to each electrode in both monkeys (Figure 1A). Both monkeys exhibited typical gross brain morphology with normal variation in the sulcal patterns in the frontal, temporal and deep brain regions where contacts where located.

### Transcranial magnetic stimulation

We conducted TMS experiments on two NHPs positioned in the sphinx posture. Both monkeys were anesthetized using Dexdomitor (0.015 mg/kg, i.m.), ketamine (6 mg/kg, i.m.), and atropine (0.045 mg/kg, i.m.), followed by 1-2% isoflurane. In each experimental session, biphasic spTMS was delivered with a MagPro X100 stimulator (MagVenture) and a butterfly coil (MC-B35, MagVenture). The coil was positioned over the left PFC for both monkeys and oriented parallel to the anterior-posterior axis. Cables and electrode connectors were kept clear of the coil and stimulator, and the amplifier was positioned well away from the monkeys’ heads during stimulation.

To study the effects of TMS pulses over the left PFC, we recorded from three implanted sEEG depth electrodes and applied active stimulation at six intensity levels: 10%, 25%, 50%, 70%, 90%, and 125% of the maximum stimulator output (MSO). To achieve 125% MSO for Monkey H pulses were delivered at 90% MSO in the stimulator’s power mode. The applied MSO intensities translated to the following current inputs (coil dI/dt): 10% MSO = 13 A/µs, 25% MSO = 37 A/µs, 50% MSO = 76 A/µs, 70% MSO = 107 A/µs, 90% MSO = 142 A/µs, and 125% MSO = 201 A/µs.

During our anesthetized recordings, we attempted to approximate the motor threshold. However, visible or electromyographic responses were difficult to elicit and highly inconsistent between trials and animals. This is in line with human and animal studies reporting markedly suppressed corticospinal excitability during anesthesia (57,58).

The monkeys were given blocks of stimulation at different intensities in a random order (Figure 1B, top), each including 80 pulses delivered at jittered inter-trial intervals of 3-5 seconds. Due to excessive motor movements induced by the pulse, only 40 trials were recorded at 125% MSO. In addition, we used a passive TMS condition as a control. In this condition the coil was rotated away from the monkey’s head to produce only the auditory click without any cortical stimulation. For the analysis presented in this paper, one block of 70% MSO passive stimulation was included for each monkey. Before each stimulation condition we recorded a baseline block (approximately one minute) without any pulses.

### Local field potentials

LFP data were acquired using a 32-channel ActiveTwo amplifier (Cortech) at a sampling rate of 40 kHz. For both monkeys, contacts in the occipital cortex (adjacent to the headpost) were chosen as the reference and ground electrodes, specifically the 9th and 11th contacts from the front of the prefrontal electrode (Figure 1A, blue electrode). To prevent the induction of electric currents at the electrode contacts due to magnetic fields, we ensured that the entire recording setup, including electrode contacts and cables, was free of loops. Additionally, the amplifier’s high input resistance (1 GV) helped minimize any induced currents to negligible levels. Data preprocessing was carried out in Python 3.11 using MNE-python (Version 1.7.1) for M/EEG analysis (59).

Preprocessing was carried out almost identically to our previous analysis (52). The raw data were segmented into epochs of 5 seconds time-locked to the TMS pulse (2500 ms pre- and post-pulse), detrended and demeaned. Noisy contacts were identified based on visual inspection and interpolated. In the next step, the TMS pulse artifact and the TMS-induced muscle artifact were linearly interpolated (−5 ms to 25 ms). Next, the LFPs were filtered using a 2^nd^-order bandstop Butterworth filter, with cutoff frequencies of 57 and 63 Hz. Finally, independent component analysis was used to decompose the signal into independent components, which were inspected and compared to the simultaneously recorded electromyography to identify and exclude residual muscle artifacts from back-projection (M_block_ = 3, SD_block_ = 1). The cleaned time series data were then resampled at 1 kHz and bandpass filtered between 0.1 Hz and 50 Hz using a 4^th^-order Butterworth filter. Examples of cleaned epochs are shown in Figure 1B (bottom).

### Time-frequency analysis

For the time-frequency analysis, all cleaned LFP data segments time-locked to pulse delivery (±2500 ms) were convolved using a complex three-cycle Morlet wavelet. The wavelet was designed to analyze 40 log-scaled frequencies from 1 Hz to 40 Hz. These wavelet parameters were chosen to replicate human EEG analyses and highlight the spectra most applicable to human EEG recordings. Total spectral power was obtained for both single trials as well as the averaged EEG spectrum across all trials for each block. Anesthesia induced through Dexdomitor and ketamine has an impact on the spectral composition of iEEG recordings which can change over time (60). This could distort results by afflicting earlier stimulation blocks differently than later blocks. In addition, potential cumulative effects of the pulses that preceded a given stimulation condition could add to this distortion. To account for the potentially time-varying effects of anesthesia and cumulative effects of TMS, total power estimates were corrected for a baseline block recorded before each stimulation condition. For each stimulation condition we randomly sampled the respective baseline block without any pulses and extracted data epochs equivalent to the ones from the stimulation (±2500 ms). These epochs were convolved using the same wavelet. The raw power estimates from the stimulation block were then divided by the average from the baseline block. Finally, relative change in the power for each condition was determined by averaging the baseline activity before a given pulse (−500 ms to -100 ms pre-pulse) across time for each frequency and then subtracting the average from each data point following stimulus presentation for the corresponding frequency. Hence, the reported power values account for possible drifts in anesthesia-related power changes and other tonic changes over time by expressing pulse-related power as a fraction of the baseline block. Further, they reflect a change in power due to the pulse by subtracting the power right before the pulse.

To ensure that no potential residual artifacts would distort the results, outlier trials were identified based on the mean power values within the first 350 ms post-pulse. If the power value at any contact at a given trial was 2.5 standard deviations above or below the mean, the trial was excluded from statistical analyses (M_block_ = 9, SD_block_ = 3). This rejection procedure resulted in an average rejection rate of 12% (SD = 3%) and the following trials counts: passive (Monkey H: 68/80, Monkey W: 72/80), 10% (Monkey H: 71/80, Monkey W: 70/80), 25% (Monkey H: 72/80, Monkey W: 68/80), 50% (Monkey H: 72/80, Monkey W: 71/80), 70% (Monkey H: 66/80, Monkey W: 72/80), 90% (Monkey H: 73/80, Monkey W: 77/80), 125% (Monkey H: 34/40).

### Statistical analysis of LFPs

Statistical analyses focused on differences between passive TMS and the varying intensities of active TMS to investigate our previous dose-dependent findings in the time-domain (52). Global differences at any electrode, time point, or frequency between TMS blocks (passive, 10%, 25%, 50%, 70%, 90%, and only for Monkey H 125%) were assessed using non-parametric cluster-level permutation tests for spatio-temporal data on single trial data from each monkey (61,62). This analysis allows for the exploration of multidimensional datasets necessitating a large number of comparisons. It addresses this issue by summarizing electrode, time point, and frequency differences into spatio-temporal clusters carrying differences between conditions. For each permutation, condition labels were randomly shuffled across trials, and an F-statistic was computed for each time-frequency-sensor sample. Samples with an F-value larger than a critical F-value were grouped to clusters of spatial and temporal adjacency based on a prior matrix reflecting the adjacency of iEEG sensors, extended by the number of frequencies and time samples. Based on these groupings, cluster-level statistics were calculated. This procedure went through 10,000 permutations to build a null distribution of the maximum cluster statistic from random partitions that cluster-level statistics were compared to. Via a Monte Carlo estimate, the permutation p-value was calculated as the proportion of random permutations yielding a larger cluster statistic than observed. To determine the critical test value, we estimated a null-hypothesis distribution and chose a conservative p-value threshold of 0.0005. Then, we calculated the critical F-value (F = 4.75) to reach significance on this level by producing an F-distribution for two-tailed testing and the corresponding degrees of freedom (N-1). Cluster permutation tests were run first for Monkey H. Then to test if they were reproducible in Monkey W the same clusters were extracted for Monkey W and F-tests equivalent to the procedure described above were run to assess differences across time-frequency and contact samples. Post-hoc analyses consisted of pairwise comparisons using two-sided independent t-tests of the averaged power values for the identified clusters. Samples were treated as independent due to the varying number of trials per condition. The resulting p-values were corrected for multiple comparisons using the FDR-method. All statistical analyses were performed with custom scripts in Python 3.11 using MNE-python (59), scipy (63), pingouin (64), and statsmodels (65).

### Phase-based connectivity analysis

Lastly, to investigate phase-based connectivity changes in response to TMS pulses, the debiased weighted phase lag index (dwPLI) calculated between the contact closest to the TMS coil (most anterior frontal contact) and all remaining contacts (66). We aimed to assess connectivity changes related to TMS pulses. Assuming TMS pulses delivered over the left PFC affect the activity of associated networks, we should find connectivity changes in relation to the seed region underneath the coil that scale with TMS intensity. A connectivity value was calculated for each time sample from 25 ms to 2000 ms after the pulse and for each frequency (1-40 Hz) resulting from the same complex Morlet wavelet transform used for our time-frequency analysis. We assume that if two regions are communicating through oscillatory changes in their LFPs, there should be a consistent phase lag between them. Hence, the dwPLI reflects the degree of asymmetry between two phase distributions, where a dwPLI of 1 indicates a consistent undirected phase lag relationship. Meanwhile, a dwPLI of 0 indicates the absence of a consistent phase lag, suggesting that phases at the two sensors occur at a random relationship. In addition, compared to the conventional phase lag index the dwPLI is more robust to confounding factors. First, it reduces the sensitivity to random noise sources by weighting the contribution of observed phase leads and lags by the magnitude of the imaginary component of the cross-spectrum. Second, it incorporates a correction for the distortion often inflicted by small sample sizes.

To assess differences between conditions, connectivity was computed for each condition by creating blocks of ten trials and calculating connectivity measures for these blocks (6-8 per condition). These were then treated as observations for statistical testing. The three time windows identified during the time-frequency analysis were considered. dwPLI values between 1-13 Hz for each of these windows were averaged across contacts with a connectivity value above 0.1 to focus on contacts relevant to the respective window. Average dwPLI values were entered as dependent variables into one-way ANOVAs with a single factor for TMS intensity comprised of one level for each intensity. Post-hoc tests were conducted using the same procedure employed for the time-frequency analysis. To ensure the homogeneity of variance and approximate normal distribution of the residuals, Levene tests and a Shapiro-Wilk tests were run. If the former indicated a violation of the sphericity assumption, the degrees of freedom were adjusted using the Greenhouse-Geisser method (67). Statistical analyses were carried out with comparable scripts to the time-frequency analysis using scipy (63), pingouin (64), and statsmodels (65) in Python 3.11.

## Results

### TMS pulses create low frequency power enhancement and suppression

To investigate the effects of spTMS over the left PFC, we recorded iEEG from two anesthetized monkeys (Monkeys H and W, Figure 1A) and applied active stimulation at six intensity levels (10%, 25%, 50%, 70%, 90%, and 125% MSO). Blocks of 80 pulses using different stimulation intensities as well as a passive condition (coil rotated away from head) were delivered in a random order (Figure 1B, top). The first analysis focused on the time-frequency convolution of the cleaned pulse-locked LFP signals (± 2500 ms), which were baseline-corrected for the time-varying influences of anesthesia. An overview of the most consistent time-frequency result patterns can be seen in Figure 2A which shows the average of all active stimulation conditions for both monkeys. For Monkey H and Monkey W there was an immediate rise in power after the pulse in a broad frequency band (∼1-20 Hz) lasting ∼400 ms, followed by a sustained suppression of power at all frequencies from approximately 400 ms to varying time points at the end of the analyzed window (approximately 1500 ms). Lastly, a rebound occurred in a subset of the contacts, with previously suppressed 1-20 Hz oscillations increasing in power. Complete time-frequency results for each monkey and all anatomical regions separated by active and passive conditions are available in the supplementary materials (Supplementary Figures S1-S4).

**Figure 2.**
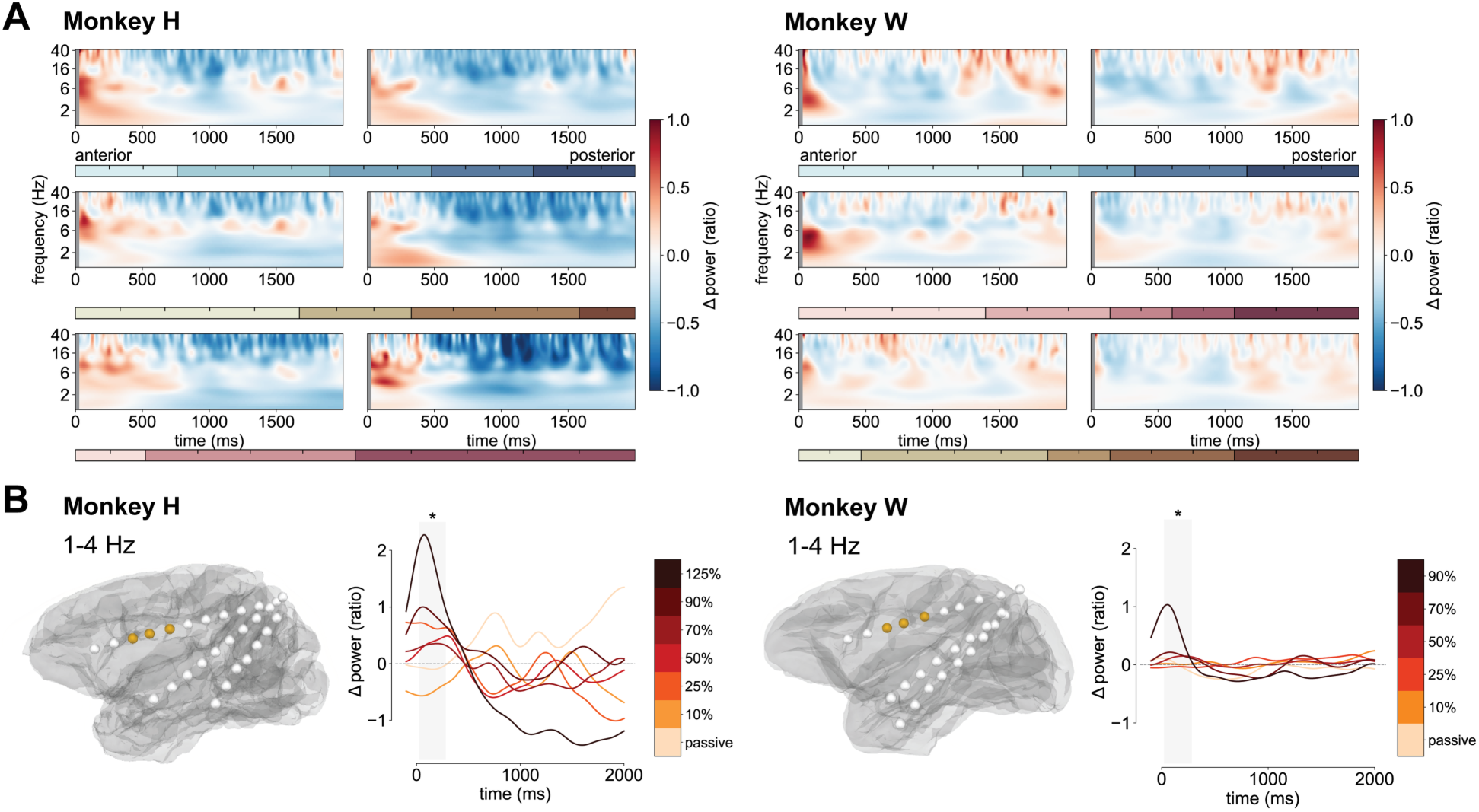
**A**: Spectrograms show the log-scaled average time-frequency response after TMS pulses for each monkey (Monkey H: left, Monkey W: right) averaged across all active stimulation intensities. Rows and horizontal color bars correspond to the electrodes shown in Figure 1A. Contacts on each electrode are split into an anterior half (left) and a posterior half (right). Gray vertical bars from 0 ms to 25 ms cover the pulse artifact corrected window. **B**: Frontal cluster between 1-4 Hz identified for Monkey H marked as gold on gray matter surfaces for both monkeys (CN, CC, ACC). Line plots to the right of each gray matter surface show the average of frequencies and contacts separated by stimulation condition. Gold to red to black colors reflect ascending stimulation intensity. * *p < 0.05*

To identify clusters of significant power modulation across conditions, we used non-parametric cluster-level permutation tests for spatio-temporal data (61,62). The analysis was first conducted on Monkey H. Subsequently, time-frequency data from the corresponding clusters were extracted for Monkey W and tested for significance (see Supplementary Table S1 for a full list of cluster details and statistics). To ensure comparability, only contacts located in the same brain regions in both monkeys were included for Monkey W. One of the first significant clusters was between 1-4 Hz and was focused on three anterior prefrontal contacts over the caudate nucleus (CN), the corpus callosum (CC), and the anterior cingulate cortex (ACC). This cluster depicted in Figure 2B was significant approximately from 25 ms post-pulse to 300 ms (Monkey H: *F_max_* = 7.18, *p* = 0.023; Monkey W: *F_max_* = 10.29, *p* = 0.033). In the later stages of the trial, a similar cluster between 1-13 Hz (25 ms to 1000 ms) emerged as significant. This cluster covered the same contacts as the first in addition to the orbitofrontal cortex (OFC), insular cortex, and the auditory core (Monkey H: *F_max_* = 8.96, *p* = 4 x 10^-4^; Monkey W: *F_max_* = 6.33, *p* = 0.037).

### Time-varying dose-dependent power modulations

Figure 3 depicts the power changes for each stimulation intensity for the 1-4 Hz and the 1-13 Hz clusters described above for both monkeys. Figure 3A illustrates the analyzed time windows and the power time courses for the passive and active conditions. There was an exponential increase in average prefrontal 1-4 Hz power across the different intensities between 25 ms and 300 ms (Figure 3B). For Monkey H passive and lower intensities (10% MSO, 50% MSO) were significantly lower in power compared to the highest intensity levels (90% MSO, 125% MSO; Supplementary Table S2). For Monkey W average 1-4 Hz power steadily increased from 10% and 25% MSO to 50% and 70% MSO, but at 90% MSO there was a sharp, exponential growth in power. Average power values for 10%, 25%, and 50% MSO were significantly lower than for 90% MSO (Supplementary Table S3).

**Figure 3.**
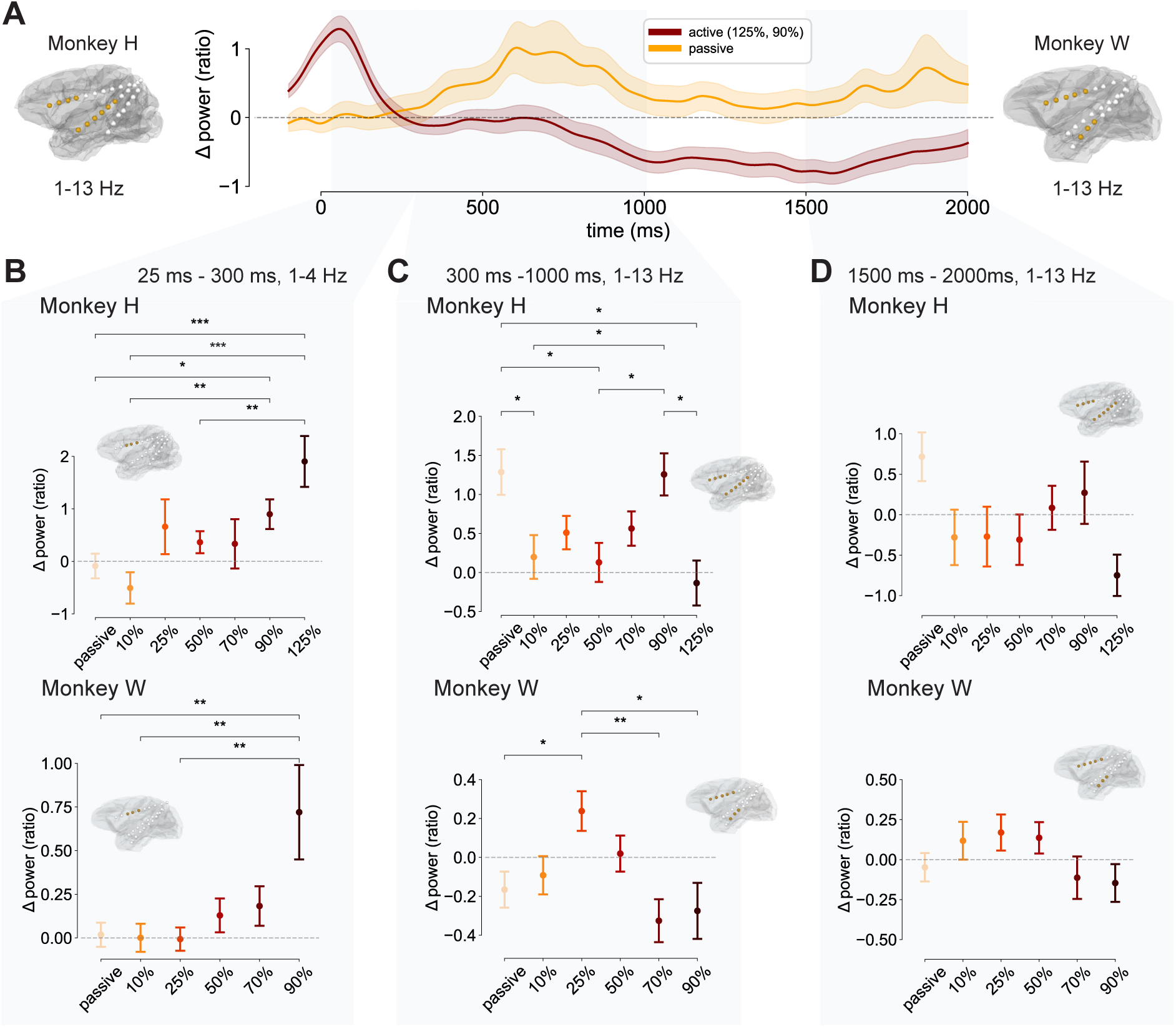
**A**: Time course of the 1-13 Hz cluster over anterior prefrontal, insular, and temporal cortical regions. Time courses are separated by active (dark red, 125% MSO for Monkey H and 90% for Monkey W) and passive stimulation (orange) averaged across monkeys. Gray overlays mark the time windows used in **B**, **C** and **D**. **B**: First cluster highlighting an early power increase immediately after the pulse delivery at the CN, CC, and ACC at 1-4 Hz between 25 ms and 300 ms. Point plots show the average power for Monkey H (top) and Monkey W (bottom) across trials per TMS pulse intensitiy. Whiskers mark the single standard error of the mean. Significant sensors included in the average are shown in gold on the respective monkey’s GM. Plots in **C** and **D** are analogous to **B** for their respective time windows (300 ms to 1000 ms for **C** and 1500 ms to 2000 ms for **D**) and contact clusters. * *p < 0.05*, ** *p < 0.01*, *** *p < 0.001*

Next, the sustained significant modulation by TMS intensity in the more broadly distributed 1-13 Hz cluster (25 ms to 1000 ms) at anterior frontal (CN, OFC, CC, ACC), medial and inferior temporal contacts (insular cortex, auditory core) was comprised of two stages (Figure 3C). After an initial rise mimicking the 1-4 Hz cluster, there was a power reduction for most active conditions (Figure 3C, 300 ms to 1000 ms). For Monkey H most active stimulation conditions were close to baseline, but power for 125% MSO was significantly lower than for the passive condition and for 90% MSO (Supplementary Table S4). A similar pattern was observed for Monkey W, with 70% MSO and 90% MSO resulting in significantly reduced power compared to 25% MSO (Supplementary Table S5). Figure 3D shows results for the end of the trial (1500 ms to 2000 ms) at the same contacts and frequencies as in Figure 3C. For both monkeys, average power in the active conditions rebounded and was closer to the level of the passive condition. Hence, there were no significant differences between conditions in this time window (Supplementary Tables S6 and S7). Notably, the average 1-13 Hz power after 125% MSO pulses for Monkey H and 90% MSO pulses for Monkey W was still negative, indicating that for these stimulation intensities power did not rebound and was still suppressed.

### Dose-dependent long- and short-range connectivity changes

Our last analysis was aimed at investigating phase-based connectivity changes in response to TMS pulses across varying intensities. For this purpose, the dwPLI was calculated between the seed contact closest to the TMS site (CN, OFC) and the remaining contacts (66). If TMS over the left PFC impacts network activity, connectivity between this seed and other contacts should change with stimulation intensity. dwPLI was calculated for every time point and frequency, using the same baseline-corrected complex Morlet wavelet transform accounting for time-varying anesthesia influences as in our time-frequency analysis. Given the results described above, analyses focused on the range of 1-13 Hz. The top panels in Figure 4 showcase the average of all active stimulation conditions for Monkey H (Figure 4A) and Monkey W (Figure 4B) separated by the time windows identified with the cluster permutation analysis run for the time-frequency analysis.

**Figure 4.**
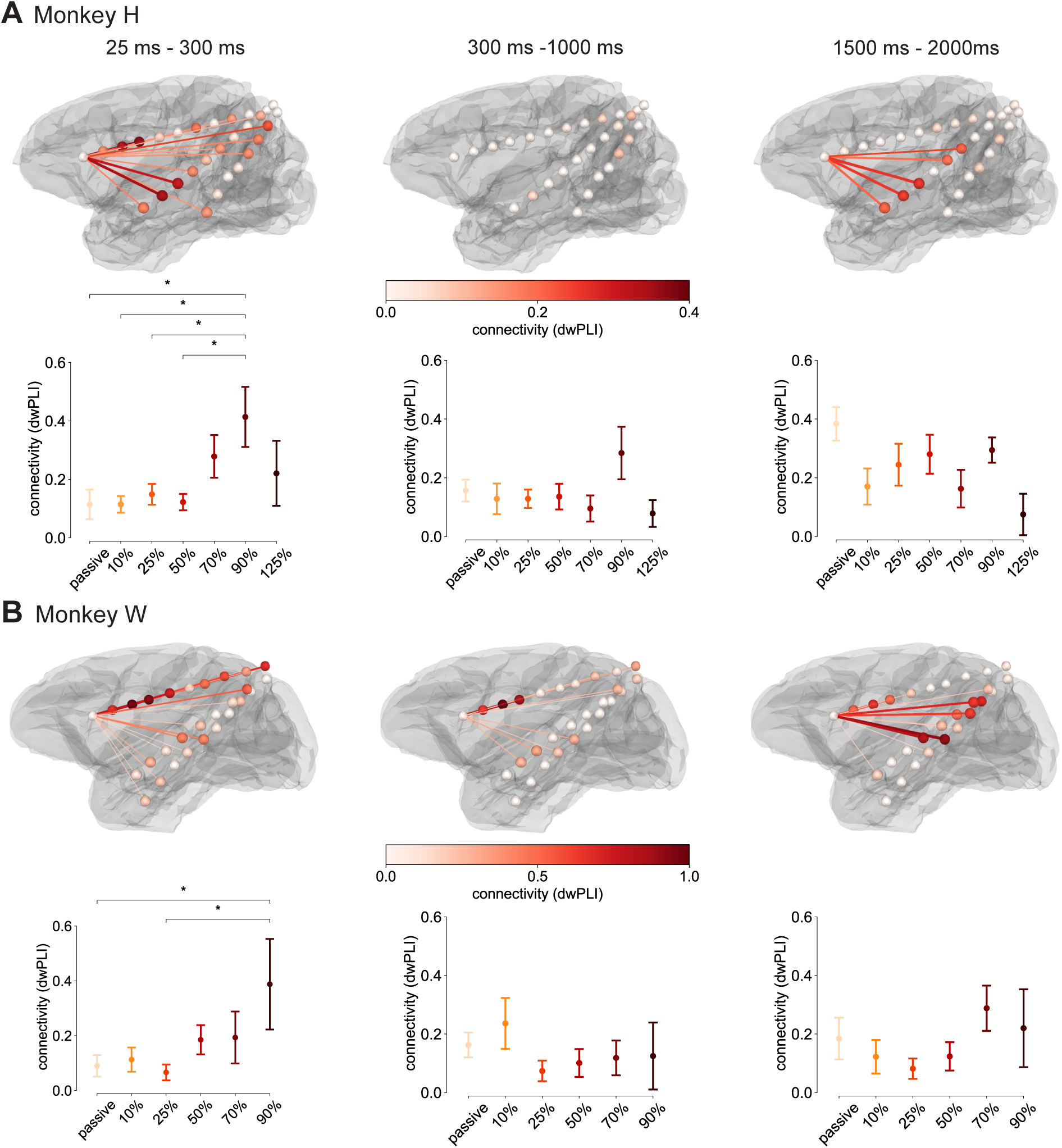
**A**: Connectivity (dwPLI) between the contact closest to the stimulation site and all remaining contacts between 1-13 Hz for the average of all active TMS conditions for Monkey H. Line colors, contact location colors, and line thickness reflect the magnitude of the connectivity estimate, with thicker lines reflecting larger connectivity. Values below 0.1 were suppressed. Columns show results for the three time windows discussed in Figure 3. Point plots depict mean connectivity values per stimulation condition averaged across contacts with a connectivity value above 0.1 separately for each time window. Since no contacts exceeded this threshold between 300 ms and 1000 ms, the same contacts as for 25 ms to 300 ms were used. Whiskers mark the single standard error of the mean. **B**: Analogous plots to **A** for Monkey W. * *p < 0.05*

Connectivity changes after active pulses generally mirrored the trends of the time-frequency analysis. First, connectivity of the stimulation site to proximal frontal contacts (CN, OFC, CC, ACC) as well as the insular cortex and the parahippocampal gyrus (PHG) increased immediately after the pulse. In addition to these contacts, posterior contacts more distal to the stimulation site (PCC, lateral intraparietal area, junction between occipital and temporal lobe) increased in connectivity. We found a significant modulation of connectivity by TMS intensity when testing it as a main effect in a one-way ANOVA (Monkey H: *F*(6, 45) = 3.37, *p* = 0.008, 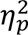 = 0.31; Monkey W: *F*(5, 34) = 2.51, *p* = 0.044, 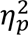 = 0.27). Post-hoc tests revealed that 90% MSO resulted in significantly larger connectivity increases relative to the passive condition (Monkey H: *t*(14) = 2.61, *p* = 0.034; Monkey W: *t*(14) = 2.22, *p* = 0.04) and 25% MSO (Monkey H: *t*(14) = 2.44, *p* = 0.036; Monkey W: *t*(14) = 2.47, *p* = 0.027). Second, approximately 300 ms after the pulse, global connectivity of the stimulation site to other contacts dropped to the level of the passive condition or below. Primarily, remaining connections from the stimulation site were to proximal regions in the CN, CC, ACC, and PCC (Figure 3B, Monkey W). However, there was no significant modulation by TMS intensity (Monkey H: *F*(6, 45) = 1.52, *p* = 0. 193, 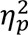 = 0.17; Monkey W: *F*(5, 34) = 1.05, *p* = 0.404, 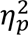 = 0.13). Third, connectivity estimates rebounded for a subset of contacts that initially increased in connectivity after the pulse, which was not significantly modulated by TMS intensity as well (Monkey H: *F*(6, 45) = 2.26, *p* = 0. 055, 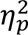 = 0.23; Monkey W: *F*(5, 34) = 1.43, *p* = 0.238, 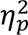 = 0.17). These contacts were in the insular cortex (Monkey H) and anterior prefrontal cortex (CN, CC, ACC) as well as the superior and medial temporal cortex (Monkey W).

## Discussion

TMS is one of the few tools to non-invasively modulate neuronal activity that has substantial clinical evidence for improving symptoms in depression (3,4). Yet, its clinical efficacy is limited by the lack of understanding of the neuronal effects of TMS, primarily due to the difficulty in monitoring neural responses to TMS in humans. To work towards a comprehensive framework for how common TMS protocols affects neuronal processing, the neuronal markers of treatment response need to be characterized using single pulses. Our study addresses this issue by investigating how spTMS applied to the left PFC modulates the oscillatory dynamics and connectivity patterns of LFPs across ipsilateral cortical regions. We consistently observed a phasic cycle of TMS pulse induced LFP changes in the range of 1-13 Hz composed of three stages: 1) An increase in frontal power close to the coil location, 2) a suppression of power across frontal, temporal, and parietal cortical regions, 3) a late power rebound at the suppressed contacts. Consistent with prior work in humans and NHPs, the effects of TMS on LFP power followed dose-dependent response curves (52,68,69). The highest intensities were coupled to higher early power enhancements and late power suppressions. Finally, we investigated phase-based connectivity changes in response to varying TMS pulses between the stimulation site and the remaining cortical contacts. Immediately following active TMS pulses, 1-13 Hz connectivity increased between the stimulation site and nearby prefrontal contacts, with additional increases observed at more distal parietal and temporal contacts relative to passive TMS. This was followed by a global connectivity reduction around 300 ms after the pulse, ending with a rebound at a subset of contacts that had initially shown increased connectivity. Only the early enhancement was modulated by TMS intensity in both monkeys, with higher intensities eliciting stronger connectivity increases.

These findings indicate that TMS creates both transient, immediate LFP changes as well as later, prolonged effects. Further, while early, transient effects seem to affect some more distant regions like the PCC and PHG, it primarily affected local regions close to and directly connected to the stimulation site such as the ventral insular cortex, the CN, and ACC. Later, sustained effects on LFP power and connectivity were spread across large-scale cortical networks including the PCC, lateral intraparietal area, as well as the superior and medial temporal gyrus. This pattern of early transient power enhancement and later sustained suppression is consistent with previous studies focusing on single cell excitation and inhibition in NHPs (50,51) and humans (70). To note, while these previous accounts discussed spiking activity patterns that unfolded on a smaller time scale (i.e., 1-50 ms), the LFP changes discussed in this paper occurred over hundreds of milliseconds or multiple seconds. This could likely be explained by the difference in neural measures (spiking vs. LFPs) and the convolution of our signals by wavelets. Importantly, while dose-dependent effects were consistent across monkeys, Monkey H was the only monkey that received 125% MSO, which achieved the highest power enhancements. However, it also resulted in the largest power reduction. This reduction was sustained until the end of the trial, which is likely the reason why 125% MSO did not exhibit a power rebound. Hence, while 125% MSO maximized power enhancements, it also suppressed LFPs and connectivity for a longer period than lower intensities.

The transient disruption followed by a return of phase synchrony is consistent with existing models of the neuronal effects of TMS. These postulate a reset of neural circuits in response to a pulse, where there is an initial wave of excitation, a transient suppression, followed by a synchronized return of activity (51,68,71). The observed dose-dependent modulation of LFP power between in the identified low frequency bands, alongside disruptions and subsequent recovery of phase-based connectivity, suggests that TMS exerts both localized and network-level effects on neural circuits. Notably, immediate power and connectivity enhancements were most pronounced at frontal contacts close to the stimulation site, but not directly underneath the coil. In addition, our connectivity results highlighted the propagation of pulse effects to distant contacts over the ventral insula, the PCC, posterior parietal cortex, and the PHG. These targets in particular fit in line with known white matter bundles connected to the lateral prefrontal cortex in humans and NHPs such as the cingulum bundle (53,72–74). This anatomical alignment reinforces the interpretation that TMS-induced activity spreads along structural pathways to downstream regions rather than through nonspecific volume conduction. Consistent with this notion, Wang et al. (2024) reported TEPs in regions directly connected to the DLPFC such as the ventral insula and the ACC when applying spTMS to the left DLPFC in human epilepsy patients. In awake NHPs oculomotor regions such as the frontal eye field and primary motor cortex have been stimulated indirectly by applying spTMS over the DLPFC (76). Moreover, rTMS protocols in NHPs applied to the left PFC have been demonstrated to reduce neuronal excitation levels and functional connectivity to wider cortical networks that overlap with the regions reported in this study including the left posterior insula, the left temporal lobe and the left postcentral gyrus (77,78). Together, these findings underscore the potential of TMS to transiently affect brain networks in a scalable manner that reflects underlying anatomical connectivity.

## Limitations

When appraising the results of this study, several limitations need to be noted. First, due the anesthesia it is possible that cortical excitability levels were reduced, leading to lower responsiveness to TMS. Second, the variability in time-frequency responses observed in Monkey H compared to Monkey W, particularly in the passive TMS condition, raises questions about individual differences in cortical excitability or anatomical factors that may influence TMS effects. The delayed low frequency power increase (∼500 ms post-pulse) observed in Monkey H likely reflected an auditory response to the TMS click rather than a direct cortical effect. This notion is supported by the fact that a few contacts in the significant cluster were located close to or within the auditory cortex. Such auditory responses may have been suppressed during active TMS due to competing stimulation effects. The stronger and more sustained response in Monkey H could stem from differences in anatomy, electrode placement, anesthesia depth, or coil positioning relative to the auditory cortex. Third, power suppression varied across intensities in both monkeys, with higher intensities more consistently producing suppression. At lower intensities, power remained elevated or near baseline, suggesting that suppression may require surpassing a threshold of initial enhancement or may scale proportionally with it. Finally, we focused our analysis on the low frequency cluster identified by the cluster permutation test due to the translational relevance of low frequency oscillations (e.g., theta) commonly studied in human EEG. Yet, the test revealed additional significant broadband clusters up to 37 Hz, which will need to be further investigated in future research.

## Conclusions

These findings emphasize the utility of NHP models for investigating the mechanisms underlying TMS. By using NHPs, the current study demonstrates how spTMS can modulate oscillatory brain activity, with significant changes in the theta band, which is thought to be particularly relevant for DLPFC functions such as cognitive control, working memory and decision-making (35,38). Furthermore, medial and lateral prefrontal theta oscillations have been suggested as markers of psychiatric disorders (36,79,80). The ability to precisely measure these effects at both local and distal cortical sites further strengthens the argument for NHP models in advancing the clinical application of TMS. Thus, this study lays a foundation for future research exploring the effects of TMS on brain oscillations and connectivity in both animal models and humans. Future studies could expand on these findings by examining the long-term effects of TMS on neural plasticity in the DLPFC, such as the potential for TMS to induce lasting changes in cortical connectivity or synaptic strength. A promising avenue for advancing TMS-based therapies is the application of closed-loop systems that integrate TMS with real-time EEG monitoring. Closed-loop TMS-EEG systems, which use real-time EEG to guide TMS delivery based on ongoing brain activity, offer a promising strategy for personalized neuromodulation by targeting specific neural states to improve treatment outcomes (81). NHP models can play a key role in refining these systems by revealing how TMS interacts with neural oscillations, helping optimize stimulation parameters and enhance clinical reliability.

