## Supplementary Materials for "Dose-dependent power and connectivity modulation of low frequency oscillations through transcranial magnetic stimulation in non-human primates"

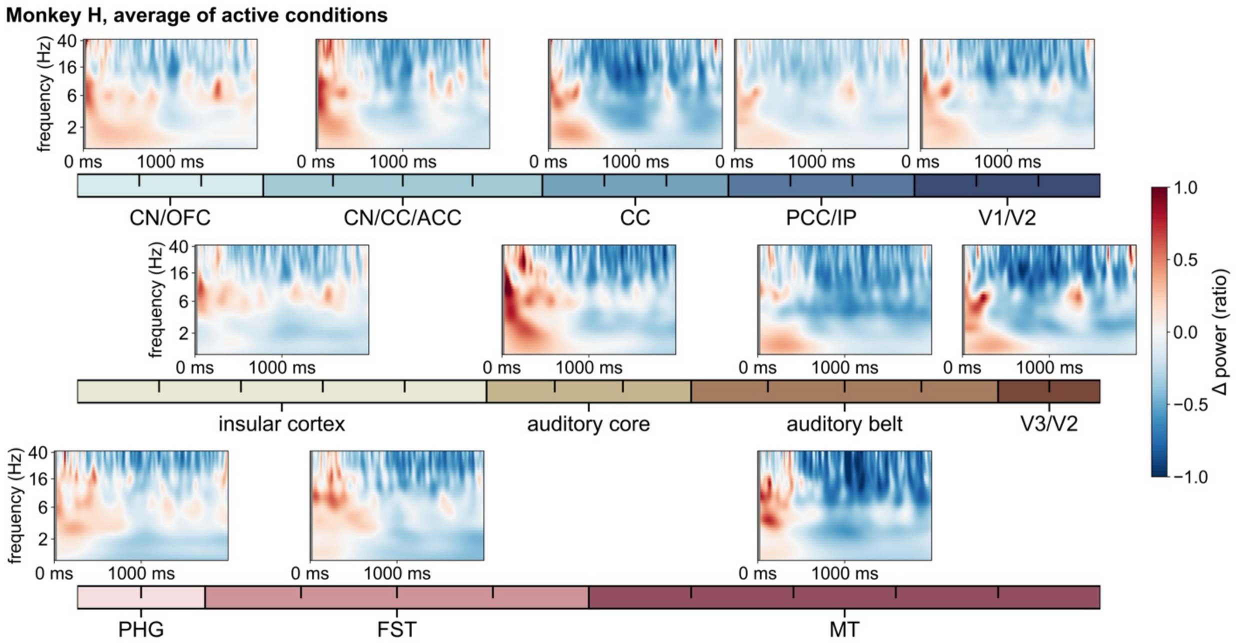


**Supplementary Figure S1.** Time-frequency results for the average of all active conditions for Monkey H. Each spectrogram represents the average of the anatomical regions listed underneath the horizontal colorbars. Grey vertical bars from 0 ms to 25 ms cover the pulse artifact corrected window.

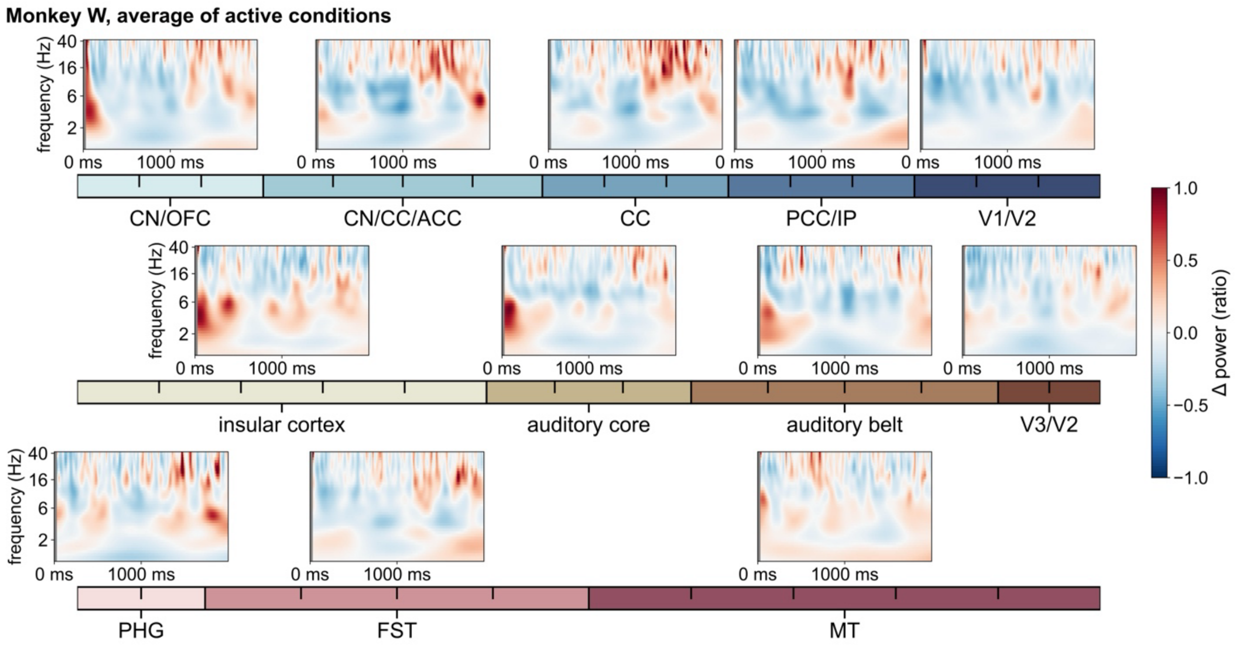


**Supplementary Figure S2.** Time-frequency results for the average of all active conditions for Monkey W. Each spectrogram represents the average of the anatomical regions listed underneath the horizontal colorbars.

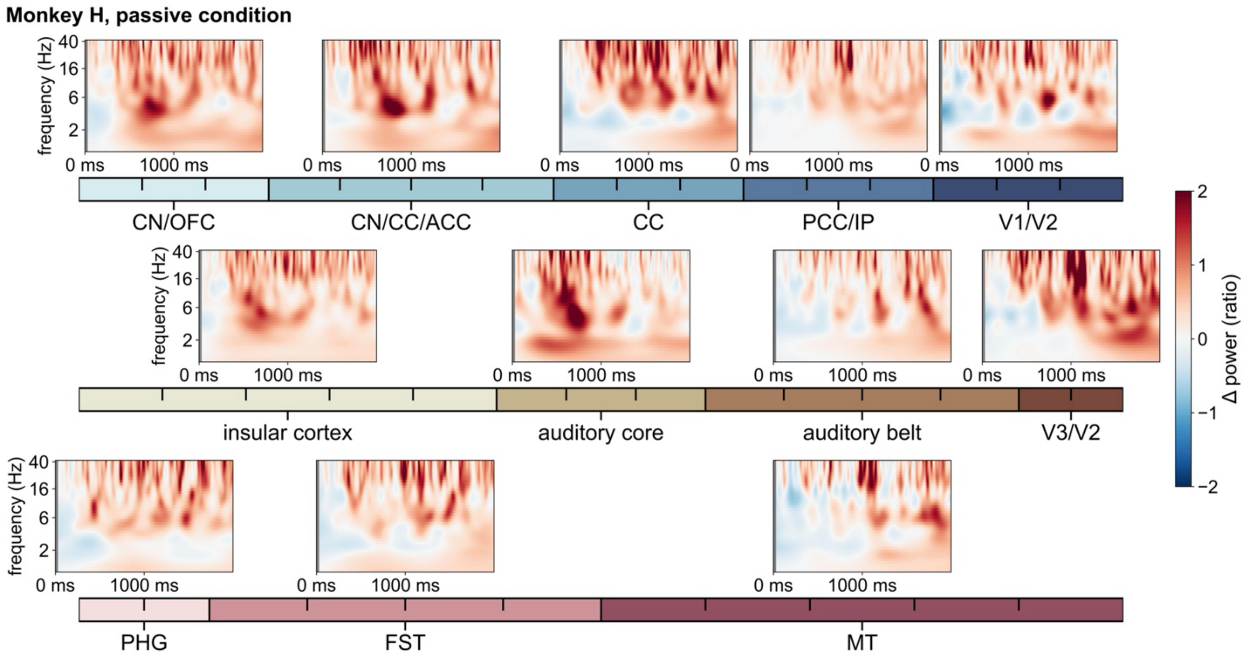


**Supplementary Figure S3.** Time-frequency results for passive condition for Monkey H. Each spectrogram represents the average of the anatomical regions listed underneath the horizontal colorbars.

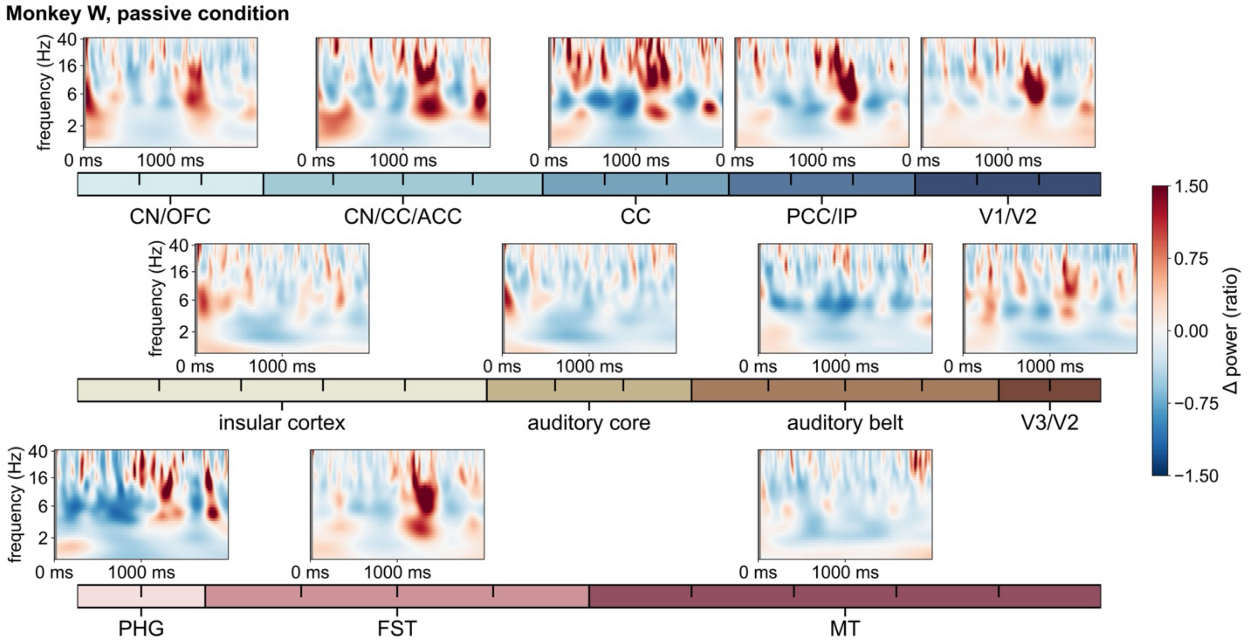


**Supplementary Figure S4.** Time-frequency results for passive condition for Monkey W. Each spectrogram represents the average of the anatomical regions listed underneath the horizontal colorbars.

**Table S1. Detailed cluster statistics for Monkey H.**

| Cluster | Time window (ms) | Contacts | Frequency range (Hz) | Cluster p-value | Max F-value |
|---|---|---|---|---|---|
| 1 | 25 - 959 | CN/OFC, CN/CC/ACC | 1 - 13 | 0.0004 | 8.961 |
| | | insular cortex, auditory core | | | |
| 2 | 25 - 287 | CN/CC/ACC | 1 - 4 | 0.0234 | 7.180 |
| 3 | 25 - 1999 | CC, PCC/IP, V1/V2 | 1 - 37 | 0.0001 | 75.475 |
| | | auditory belt, V3/V2 | | | |
| | | FST, MT | | | |
| 4 | 25 - 275 | CN/OFC, CN/CC/ACC, CC | 4 - 37 | 0.0015 | 35.670 |
| | | insular cortex, auditory core | | | |
| 5 | 168 - 218 | PCC/IP | 19 - 37 | 0.0434 | 113.674 |
| 6 | 263 - 319 | PCC/IP | 13 - 37 | 0.0353 | 68.897 |
| 7 | 264 - 312 | auditory belt | 17 - 37 | 0.0309 | 89.737 |
| 8 | 760 - 830 | PCC/IP | 11 - 28 | 0.0361 | 55.801 |
| 9 | 760 - 831 | auditory belt | 11 - 28 | 0.0474 | 48.989 |
| 10 | 967 - 1123 | CN/OFC, CN/CC/ACC, CC, PCC/IP, V1/V2 | 12 - 37 | 0.0061 | 15.885 |
| | | insular cortex, auditory core, auditory belt, V3/V2 | | | |
| | | FST | | | |
| 11 | 1452 - 1719 | auditory belt | 3 - 7 | 0.0306 | 14.316 |
| 12 | 1461 - 1675 | PCC/IP | 3 - 10 | 0.0376 | 11.975 |
| 13 | 1740 - 1895 | auditory belt | 9 - 18 | 0.0368 | 25.786 |
| 14 | 1897 - 1999 | PCC/IP, V1/V2 | 12 - 37 | 0.0302 | 18.431 |
| | | auditory belt | | | |

**Supplementary Table S1.** List of significant clusters ($p < 0.05$) for Monkey H. Columns show details about time points, frequency points and contacts included in the cluster as well as cluster-level statistics. Contact labels are the anatomical regions contacts were located in. Rows nested within a cluster correspond to the three electrodes contacts were located on. Cluster-level p-values are rounded to the second position behind the decimal point. F-values are the maximum F-value in a given cluster. CN: caudate nucleus, OFC: orbitofrontal cortex, CC: corpus callosum, ACC: anterior cingulate cortex, PCC: posterior cingulate cortex, IP: intraparietal area, V1: visual area 1, V2: visual area 2, V3: visual area 3, PHG: parahippocampal gyrus, FST: fundus of the superior temporal visual area, MT: middle temporal area.

**Table S2. Pairwise comparisons (Monkey H, 1-4 Hz, 25-300 ms, CN, CC, ACC).**

| Condition 1 | Condition 2 | Mean difference | SEM | t-value | p-value | Significance |
|---|---|---|---|---|---|---|
| 10% | passive | -0.42 | 0.38 | -1.11 | 0.370 | |
| 25% | passive | 0.75 | 0.57 | 1.29 | 0.298 | |
| 50% | passive | 0.45 | 0.32 | 1.44 | 0.247 | |
| 70% | passive | 0.42 | 0.52 | 0.85 | 0.489 | |
| **90%** | **passive** | **0.99** | **0.37** | **2.68** | **0.035** | * |
| **125%** | **passive** | **1.99** | **0.54** | **4.17** | **< 0.001** | *** |
| 25% | 10% | 1.17 | 0.60 | 1.88 | 0.145 | |
| 50% | 10% | 0.87 | 0.36 | 2.43 | 0.057 | |
| 70% | 10% | 0.84 | 0.56 | 1.55 | 0.246 | |
| **90%** | **10%** | **1.41** | **0.41** | **3.43** | **0.005** | ** |
| **125%** | **10%** | **2.41** | **0.57** | **4.45** | **< 0.001** | *** |
| 50% | 25% | -0.30 | 0.56 | -0.53 | 0.696 | |
| 70% | 25% | -0.33 | 0.70 | -0.45 | 0.721 | |
| 90% | 25% | 0.24 | 0.59 | 0.40 | 0.723 | |
| 125% | 25% | 1.24 | 0.71 | 1.49 | 0.246 | |
| 70% | 50% | -0.03 | 0.51 | -0.06 | 0.949 | |
| 90% | 50% | 0.53 | 0.35 | 1.53 | 0.246 | |
| **125%** | **50%** | **1.54** | **0.53** | **3.41** | **0.005** | ** |
| 90% | 70% | 0.56 | 0.55 | 1.08 | 0.370 | |
| 125% | 70% | 1.57 | 0.68 | 2.20 | 0.093 | |
| 125% | 90% | 1.00 | 0.56 | 1.90 | 0.145 | |

**Supplementary Table S2.** Pairwise comparisons using independent t-tests for Monkey H and the 1-4 Hz cluster at prefrontal regions (CN: Caudate Nucleus, CC: Corpus Collosum, ACC: Anterior Cingulate Cortex) between 25 ms and 300 ms. Condition columns show the pairing of conditions being compared in a given row. Statsictis for the respective pair including their mean power difference (ratio change), the standard error of the mean (SEM), t-value, and corrected p-value are shown to the right. All p-values were corrected using the FDR method. Significance is marked by bold text as well as the notation in the significance colum: * $p < 0.05$, ** $p < 0.01$, *** $p < 0.001$.

**Table S3. Pairwise comparisons (Monkey W, 1-4 Hz, 25-300 ms, CN, CC, ACC).**

| Condition 1 | Condition 2 | Mean difference | SEM | t-value | p-value | Significance |
|---|---|---|---|---|---|---|
| 10% | passive | -0.02 | 0.11 | -0.16 | 0.932 | |
| 25% | passive | -0.03 | 0.10 | -0.26 | 0.917 | |
| 50% | passive | 0.11 | 0.12 | 0.95 | 0.467 | |
| 70% | passive | 0.16 | 0.13 | 1.28 | 0.380 | |
| **90%** | **passive** | **0.70** | **0.28** | **3.50** | **0.006** | ** |
| 25% | 10% | -0.01 | 0.10 | -0.07 | 0.941 | |
| 50% | 10% | 0.13 | 0.13 | 1.02 | 0.464 | |
| 70% | 10% | 0.18 | 0.14 | 1.29 | 0.380 | |
| **90%** | **10%** | **0.72** | **0.28** | **3.25** | **0.009** | ** |
| 50% | 25% | 0.14 | 0.12 | 1.20 | 0.390 | |
| 70% | 25% | 0.19 | 0.13 | 1.50 | 0.342 | |
| **90%** | **25%** | **0.73** | **0.28** | **3.67** | **0.006** | ** |
| 70% | 50% | 0.05 | 0.15 | 0.36 | 0.903 | |
| 90% | 50% | 0.59 | 0.29 | 2.51 | 0.054 | |
| 90% | 70% | 0.54 | 0.29 | 2.17 | 0.100 | |

**Supplementary Table S3.** Pairwise comparisons using independent t-tests for Monkey W and the 1-4 Hz cluster at prefrontal regions regions between 25 ms and 300 ms. The table is structured identically to Supplementary Table S2. ** $p < 0.01$.

**Table S4. Pairwise comparisons (Monkey H, 1-13 Hz, 300-1000 ms, CN, CC, ACC, insular cortex, auditory core).**

| Condition 1 | Condition 2 | Mean difference | SEM | t-value | p-value | Significance |
|---|---|---|---|---|---|---|
| **10%** | **passive** | **-1.09** | **0.40** | **-2.68** | **0.038** | ***** |
| 25% | passive | -0.78 | 0.36 | -2.08 | 0.103 | |
| **50%** | **passive** | **-1.16** | **0.38** | **-2.78** | **0.038** | ***** |
| 70% | passive | -0.72 | 0.37 | -1.91 | 0.122 | |
| 90% | passive | -0.03 | 0.40 | -0.07 | 0.944 | |
| **125%** | **passive** | **-1.42** | **0.41** | **-2.53** | **0.046** | ***** |
| 25% | 10% | 0.31 | 0.35 | 0.88 | 0.499 | |
| 50% | 10% | -0.07 | 0.37 | -0.17 | 0.905 | |
| 70% | 10% | 0.37 | 0.36 | 1.02 | 0.436 | |
| **90%** | **10%** | **1.06** | **0.39** | **2.65** | **0.038** | ***** |
| 125% | 10% | -0.33 | 0.40 | -0.64 | 0.617 | |
| 50% | 25% | -0.38 | 0.33 | -1.16 | 0.373 | |
| 70% | 25% | 0.05 | 0.31 | 0.17 | 0.905 | |
| 90% | 25% | 0.75 | 0.34 | 2.20 | 0.090 | |
| 125% | 25% | -0.65 | 0.36 | -1.60 | 0.197 | |
| 70% | 50% | 0.43 | 0.33 | 1.31 | 0.314 | |
| **90%** | **50%** | **1.13** | **0.37** | **3.07** | **0.030** | ***** |
| 125% | 50% | -0.26 | 0.38 | -0.63 | 0.617 | |
| 90% | 70% | 0.69 | 0.35 | 2.02 | 0.108 | |
| 125% | 70% | -0.70 | 0.36 | -1.71 | 0.172 | |
| **125%** | **90%** | **-1.39** | **0.39** | **-3.10** | **0.030** | ***** |

**Supplementary Table S4.** Pairwise comparisons using independent t-tests for Monkey H and the 1-13 Hz cluster at prefrontal and temporal regions between 300 ms and 1000 ms. The table is structured identically to Supplementary Table S2. * $p < 0.05$.

**Table S5. Pairwise comparisons (Monkey W, 1-13 Hz, 300-1000 ms, CN, CC, ACC, insular cortex, auditory core).**

| Condition 1 | Condition 2 | Mean difference | SEM | t-value | p-value | Significance |
|---|---|---|---|---|---|---|
| 10% | passive | 0.07 | 0.14 | 0.55 | 0.626 | |
| **25%** | **passive** | **0.40** | **0.14** | **2.90** | **0.032** | * |
| 50% | passive | 0.19 | 0.13 | 1.41 | 0.267 | |
| 70% | passive | -0.16 | 0.14 | -1.12 | 0.398 | |
| 90% | passive | -0.11 | 0.17 | -0.63 | 0.608 | |
| 25% | 10% | 0.33 | 0.14 | 2.30 | 0.069 | |
| 50% | 10% | 0.11 | 0.13 | 0.82 | 0.515 | |
| 70% | 10% | -0.23 | 0.15 | -1.59 | 0.224 | |
| 90% | 10% | -0.18 | 0.17 | -1.04 | 0.411 | |
| 50% | 25% | -0.22 | 0.14 | -1.57 | 0.224 | |
| **70%** | **25%** | **-0.56** | **0.15** | **-3.71** | **0.005** | ** |
| **90%** | **25%** | **-0.51** | **0.18** | **-2.70** | **0.040** | * |
| 70% | 50% | -0.34 | 0.14 | -2.42 | 0.064 | |
| 90% | 50% | -0.29 | 0.17 | -1.73 | 0.217 | |
| 90% | 70% | 0.05 | 0.18 | 0.27 | 0.786 | |

**Supplementary Table S5.** Pairwise comparisons using independent t-tests for Monkey W and the 1-13 Hz cluster at prefrontal and temporal regions between 300 ms and 1000 ms. The table is structured identically to Supplementary Table S2. * $p < 0.05$, ** $p < 0.01$.

**Table S6. Pairwise comparisons (Monkey H, 1-13 Hz, 1500-2000 ms, CN, CC, ACC, insular cortex, auditory core).**

| Condition 1 | Condition 2 | Mean difference | SEM | t-value | p-value | Significance |
|---|---|---|---|---|---|---|
| 10% | passive | -1.00 | 0.46 | -2.18 | 0.209 | |
| 25% | passive | -0.99 | 0.48 | -2.07 | 0.209 | |
| 50% | passive | -1.03 | 0.43 | -2.36 | 0.203 | |
| 70% | passive | -0.63 | 0.41 | -1.56 | 0.365 | |
| 90% | passive | -0.45 | 0.49 | -0.91 | 0.543 | |
| 125% | passive | -1.47 | 0.39 | -2.94 | 0.083 | |
| 25% | 10% | 0.01 | 0.50 | 0.02 | 0.985 | |
| 50% | 10% | -0.03 | 0.46 | -0.06 | 0.985 | |
| 70% | 10% | 0.36 | 0.44 | 0.83 | 0.543 | |
| 90% | 10% | 0.55 | 0.51 | 1.07 | 0.543 | |
| 125% | 10% | -0.47 | 0.43 | -0.84 | 0.543 | |
| 50% | 25% | -0.04 | 0.48 | -0.08 | 0.985 | |
| 70% | 25% | 0.35 | 0.46 | 0.78 | 0.543 | |
| 90% | 25% | 0.54 | 0.53 | 1.02 | 0.543 | |
| 125% | 25% | -0.48 | 0.45 | -0.80 | 0.543 | |
| 70% | 50% | 0.39 | 0.41 | 0.95 | 0.543 | |
| 90% | 50% | 0.58 | 0.50 | 1.17 | 0.543 | |
| 125% | 50% | -0.44 | 0.40 | -0.86 | 0.543 | |
| 90% | 70% | 0.19 | 0.47 | 0.40 | 0.808 | |
| 125% | 70% | -0.83 | 0.37 | -1.83 | 0.291 | |
| 125% | 90% | -1.02 | 0.46 | -1.64 | 0.364 | |

**Supplementary Table S6.** Pairwise comparisons using independent t-tests for Monkey H and the 1-13 Hz cluster at prefrontal and temporal regions between 1500 ms and 2000 ms. The table is structured identically to Supplementary Table S2.

**Table S7. Pairwise comparisons (Monkey W, 1-13 Hz, 1500-2000 ms, CN, CC, ACC, insular cortex, auditory core).**

| Condition 1 | Condition 2 | Mean difference | SEM | t-value | p-value | Significance |
|---|---|---|---|---|---|---|
| 10% | passive | 0.17 | 0.15 | 1.13 | 0.431 | |
| 25% | passive | 0.22 | 0.14 | 1.49 | 0.365 | |
| 50% | passive | 0.18 | 0.13 | 1.40 | 0.365 | |
| 70% | passive | -0.07 | 0.16 | -0.42 | 0.904 | |
| 90% | passive | -0.10 | 0.15 | -0.62 | 0.801 | |
| 25% | 10% | 0.05 | 0.16 | 0.31 | 0.904 | |
| 50% | 10% | 0.02 | 0.15 | 0.12 | 0.904 | |
| 70% | 10% | -0.23 | 0.18 | -1.30 | 0.365 | |
| 90% | 10% | -0.26 | 0.17 | -1.36 | 0.365 | |
| 50% | 25% | -0.03 | 0.15 | -0.22 | 0.904 | |
| 70% | 25% | -0.28 | 0.17 | -1.62 | 0.365 | |
| 90% | 25% | -0.32 | 0.16 | -1.58 | 0.365 | |
| 70% | 50% | -0.25 | 0.17 | -1.54 | 0.365 | |
| 90% | 50% | -0.28 | 0.15 | -1.68 | 0.365 | |
| 90% | 70% | -0.03 | 0.18 | -0.16 | 0.904 | |

**Supplementary Table S7.** Pairwise comparisons using independent t-tests for Monkey W and the 1-13 Hz cluster at prefrontal and temporal regions between 1500 ms and 2000 ms. The table is structured identically to Supplementary Table S2.
